# Exercise and training regulation of autophagy markers in human and rodent skeletal muscle

**DOI:** 10.1101/2021.04.06.437625

**Authors:** Javier Botella, Nicholas A. Jamnick, Cesare Granata, Amanda J. Genders, Enrico Perri, Tamim Jabar, Andrew Garnham, Michael Lazarou, David J. Bishop

**Affiliations:** Institute for Health and Sport (iHeS), Victoria University, Melbourne, Victoria, Australia; Metabolic Research Unit, School of Medicine and Institute for Mental and Physical Health and Clinical Translation (IMPACT), Deakin University, Waurn Ponds, Victoria, Australia; Department of Diabetes, Central Clinical School, Monash University, Melbourne, Victoria, Australia; Department of Biomedical Sciences for Health, University of Milan, Italy; Department of Biochemistry and Molecular Biology, Biomedicine Discovery Institute, Monash University, Melbourne, Australia

**Keywords:** autophagy, exercise, LC3, skeletal muscle

## Abstract

Autophagy is a key intracellular mechanism by which cells degrade old or dysfunctional proteins and organelles. In skeletal muscle, evidence suggests that exercise increases autophagosome content and autophagy flux. However, the exercise-induced response seems to differ between rodents and humans, and little is known about how different exercise prescription parameters may affect these results. The present study utilised skeletal muscle samples obtained from four different experimental studies using rats and humans. Here we show that following exercise, in the soleus muscle of Wistar rats, there is an increase in LC3B-I protein levels (+ 109%) immediately after exercise, and a subsequent increase in LC3B-II protein levels (+ 97%) 3 hours into the recovery. Conversely, in human skeletal muscle, there is an immediate exercise-induced decrease in LC3B-II protein levels (− 24%), independent of whether exercise is performed below or above the maximal lactate steady state, which returns to baseline 3.5 hours following recovery, while no change in LC3B-I protein levels is observed. p62 protein levels did not change in neither rats nor humans following exercise. By employing an *ex vivo* autophagy flux assay previously used in rodents we demonstrate that the exercise-induced decrease in LC3B-II protein levels in humans does not reflect a decreased autophagy flux. Instead, effect size analyses suggest a modest-to-large increase in autophagy flux following exercise that lasts up to 24 hours. Our findings suggest that exercise-induced changes in autophagosome content markers differ between rodents and humans, and that exercise-induced decrease in LC3B-II protein levels do not reflect autophagy flux level.

## Introduction

Autophagy is the cellular process by which an autophagosome (a double–membrane vesicle) engulfs, and delivers to the lysosome, proteins or organelles that need to be degraded. It is the recycling machinery of the cell and is important for the correct removal of intracellular pathogens or misfolded proteins, among others, which may activate deleterious cellular signalling pathways (e.g., inflammation) (1). In skeletal muscle, autophagy is important to prevent mitochondrial damage (2), as well as promoting positive muscle regeneration (3), optimal glucose metabolism (4), and for training-induced increases in mitochondrial proteins and endurance performance (5). Thus, it is important to better understand the factors that influence autophagy in skeletal muscle.

The autophagy machinery consists of a core set of autophagy-related (ATG) proteins (6). Among these, the ATG8 family (which includes the subfamily members LC3A, LC3B, LC3C, GABARAP, GABARAPL1, GABARAPL2) promote autophagosome formation and autophagosome-lysosome fusion (7, 8). In skeletal muscle LC3s are abundantly expressed (9), which makes LC3 a widely used marker of autophagosome content. In rodents, endurance exercise acutely increases the levels of LC3-II (and the LC3-II/I ratio) as well as the appearance of LC3 puncta in both skeletal and cardiac muscle (4). Similarly, endurance exercise to exhaustion increases LC3-II protein levels in the tibialis anterior, along with a tendency for increased exercise-induced autophagy flux, but unchanged protein levels of the autophagy receptor p62 (10). Following a similar exercise session in mice LC3-I protein levels have also been reported to increase (11). A study comparing two different exercise regimes in mice showed that both low- and moderate-intensity exercise increased LC3A/B-II protein levels and the LC3A/B-II/I ratio 3 hours following the end of exercise (12). Although findings are inconclusive with regard to exercise-induced p62 protein changes, the LC3 findings collectively suggest that autophagosome content, and possibly autophagy flux, are increased after endurance exercise in rodents.

In contrast to rodent studies, human studies show a distinct pattern of exercise-induced changes in autophagosome content markers. Protein levels of LC3B-II, and the LC3B-II/I ratio, have been shown to decrease 0 to1 h following different types of endurance exercise and return to baseline values after 3 to 4 h of recovery in human skeletal muscle (13–16). In contrast, 60 min of exercise at 60% of V̇O_2max_ has been reported to increase the levels of LC3A/B-II protein levels 2 h after the end of exercise (17). Following most types of endurance exercise, the protein levels of the autophagy receptor p62 remained unchanged (13, 15–17). In contrast, following 2 h at 70% of V̇O_2peak_, but not at 55% of V̇O_2peak_, the p62 protein levels decreased, which could suggest an effect of exercise intensity on the exercise-induced p62 protein changes (14). However, since both the exercise intensity and the total work completed were different, it is difficult to isolate any of these factors. Other differences such as training status, timing of biopsies, antibodies used, or sample size may also contribute to the reported discrepancies between human studies. Whether exercise intensity distinctly affects the LC3B and p62 protein levels changes following an exercise session remains to be fully elucidated.

Autophagy flux assays are considered the ‘gold-standard’ to assess autophagy levels (18). Autophagy flux is the term used for the combined autophagy steps, which includes autophagosome formation, maturation, fusion with lysosomes, and breakdown of the autolysosome contents. One such assay aims to chemically block the fusion of autophagosomes with the lysosome (the end-point of the degradation process) and to monitor the accumulation of LC3-II (18). Performing an *in vivo* autophagy flux is not ethically possible in human tissues and remains a limitation. This means that human studies have relied on markers of autophagosome and autophagy receptor protein levels (13–17). Although not previously used in humans, animal models have also utilised an *ex vivo* autophagy flux analysis (19). Implementing this *ex vivo* LC3-II flux assay could provide a direct assessment of autophagy in human studies and would avoid having to rely solely on indirect markers (i.e., LC3-II/I ratio).

Despite the increase in autophagy research in skeletal muscle, there is currently no consensus on the exercise-induced regulation of autophagosome content in skeletal muscle. The aims of the current study are multiple: 1) to assess potential differences in exercise-induced changes in LC3B and p62 protein between rodents and humans; 2) to elucidate if the exercise-induced LC3B and p62 protein changes are affected by exercise below or above the maximal lactate steady state (MLSS) in humans; 3) to explore the effects of exercise training on the basal LC3B and p62 protein levels in humans; and 4) to assess whether the exercise-induced changes in LC3B-II protein levels are reflective of a decreased *ex vivo* autophagy flux in humans.

## Materials and Methods

Four different studies were included in this manuscript: a single exercise session in rats (Study 1), exercise in humans at three different work-matched intensities above or below the maximal lactate steady state (MLSS) (Study 2), a 3-week high-volume high-intensity interval training in humans (Study 3), and a single exercise session in humans for the establishment of *ex vivo* autophagy flux (Study 4). All human participants were deemed healthy, and their characteristics can be found in Table 1. Studies were performed at Victoria University (Melbourne, Australia), and all analyses were performed under similar conditions in the same laboratory. All studies were approved by the Victoria University Animal Ethics Committee and the Victoria University Human Research Ethics Committee. Informed consent was obtained from all human participants prior to study participation.

**Table 1.** Descriptive data of the human participants recruited for studies 2, 3, and 4. Data are mean ± SD.

| | Age<br>(y) | $\dot{V}O_{2peak}$<br>(mL·min<sup>-1</sup>·kg<sup>-1</sup>) | Trial | Relative<br>Exercise<br>Intensity<br>(% $\dot{W}_{max}$) | Absolute<br>Exercise Intensity<br>(W) |
| --- | --- | --- | --- | --- | --- |
| Study 2<br>(n = 10) | 27.5<br>± 7.7 | 55.8<br>± 10.0 | - 18 % MLSS | 51 ± 4 | 181 ± 39 |
| | | | - 6 % MLSS | 59 ± 4 | 207 ± 45 |
| | | | + 6 % MLSS | 67 ± 5 | 234 ± 51 |
| Study 3<br>(n = 9) | 22.4<br>± 5.2 | 47.0<br>± 7.5 | - | | |
| Study 4<br>(n = 5) | 30.0<br>± 7.3 | 48.1<br>± 4.4 | SIE | 175 ± 21 | 444 ± 179 |
| | | | MICE | 42 ± 2 | 142 ± 35 |

### Study 1 – Exercise in rats

#### Overview

Twenty-eight male Wistar rats (8 weeks old) were obtained from the Animal Resource Centre (Perth, Australia). The Victoria University Animal Ethics Committee approved this study (AEC 15/002). All procedures were performed according to the Australian Code of Practice for the Care and Use of Animals for Scientific Purposes (National Health and Medical Research Council, Australia, 8th Edition). Rats were housed in groups of 2 to 4 in a temperature-controlled room and maintained with a chow diet (Specialty Feeds, Perth, WA) and water ad libitum on a 12:12 h light-dark cycle, 18-22 °C, with approximately 50% humidity. The animals underwent acclimatisation over three days, using five separate 15 min running sessions (ranging from being placed on a non-moving treadmill belt to running at a speed of 0.25 m·s^−1^). At least 48 h before the experimental exercise session, the animals performed an incremental exercise test. The incline of the treadmill was set at 10 degrees and the test was started at 0.16 m·s^−1^. The speed of the treadmill was increased 0.05 m·s^−1^ every three minutes. Animals were removed from the treadmill when they could no longer keep up with the speed despite encouragement (air puff).

#### Experimental session

On the experimental day, animals were exercised at 80% of their top speed achieved during the incremental test (approximately 0.38 m·s^−1^ at a 10 degree incline) for seven 2-minute intervals interspersed with 1 minute of rest. Rats were humanely killed using 90 mg·kg^−1^ i.p. pentobarbitone prior to (REST), immediately after (+0 h), or 3 h after the completion of the exercise protocol, and the soleus was removed and immediately frozen in liquid nitrogen and stored at −80 °C.

### Study 2 – Exercise in humans: effects of exercise intensity

#### Overview

Ten healthy males volunteered for this study (Table 1). Participants were required to attend the laboratory at Victoria University 8 to 11 times. For the first trial, participants underwent a cycling graded exercise test (GXT) with 1-min increments as previously described (20). The following visits were dedicated to determining the maximal lactate steady state (MLSS), which was established by a series of 30-min constant power sessions. After the establishment of the MLSS, participants completed two constant power exercise sessions to exhaustion at +6% of the MLSS. Following this, they performed three experimental sessions that included skeletal muscle biopsies.

#### Experimental session

The three experimental sessions were performed in a randomised order at −18%, −6% or +6% of the MLSS. The MLSS was selected as the reference point because it is a critical intensity that delineates heavy from severe exercise intensity (21), and three intensities (2 below and 1 above the MLSS) were chosen for the study. Participants were given 48 h of complete rest before each trial, and at least 7 days between the successive experimental trials. They were asked to maintain their normal diet and to replicate it on the day before and during the experimental trials. Biopsies were taken from the *vastus lateralis* muscle at rest before the start of exercise (REST), immediately upon completion of the exercise session (+ 0 h), and 3.5 h after the end of the exercise (+ 3.5 h). Samples were immediately cleaned of excess blood, fat, or connective tissue, and rapidly frozen in liquid nitrogen. Samples were stored at −80 °C until subsequent analyses.

### Study 3 - Exercise training in humans: effect of high-volume training

#### Overview

Participants completed 20 days of twice-a-day high-intensity interval training (HIIT), as previously published (22, 23). Skeletal muscle biopsies were obtained at rest before (PRE) and after (POST) the 20 days of high-volume of HIIT.

#### Experimental sessions

Participants were given 48 h of rest before the sample collection. All samples were obtained from the *vastus lateralis* muscle and participants were provided standardised meals, as previously described (23, 24). Biopsies were taken at rest and were immediately cleaned of excess blood, fat, or connective tissue, and rapidly frozen in liquid nitrogen and stored at −80 °C for subsequent analyses.

### Study 4 – Exercise-induced autophagy flux in human skeletal muscle

#### Overview

Samples from five healthy participants from a larger study were analysed. The GXT protocol utilised in this study was the same as in Study 2. Participants had been familiarised with the exercise required as they had undertaken two GXTs and two exercise sessions in the two weeks before the experimental session.

#### Experimental session

Two participants underwent the following exercise: six 30-s ‘all-out’ cycling bouts against a resistance initially set at 0.075 kg·kg body mass^−1^ (~ 175% Ẇ_max_), interspersed with a 4-min recovery period. The other three participants performed a session consisting of 90 minutes of continuous cycling at ~ 42% of Ẇ_max_. Participants were given 72 h of rest before the experimental session. All samples were obtained from the *vastus lateralis* muscle and participants were provided standardised meals, as in previous studies (23, 24). Biopsies were taken at rest before the start of exercise (REST), immediately upon completion of the exercise bout (+ 0 h), 2.5 hours into the recovery (+ 2.5 h), and 24 hours after the initial skeletal muscle sample (+ 24 h). Small muscle portions were immediately immersed into two separate vials with 3 mL of oxygenated DMEM, and the autophagy flux assay was started (see below in *ex vivo* autophagy flux assay). Once the protocol was finalised, samples were stored at −80 °C for subsequent analyses

#### Skeletal Muscle Analyses

##### Preparation of whole-muscle lysates

Approximately 10 to 20 mg of frozen muscle was homogenised two times for two minutes at a speed of 30 Hz with a TyssueLyser instrument (Qiagen, Canada) in an ice-cold lysis buffer (1:20 w/v) containing 50 mM Tris-HCl, 150 mM NaCl, 1 mM EDTA, 5 mM Na_4_P_2_0_7_, 1 mM Na_3_V0_4_, 1 % NP-40, with added protease and phosphatase inhibitors at a 1:100 concentration (Cell Signaling Technology). Protein concentration was determined using a commercial colourimetric assay (Bio-Rad Protein Assay kit II, Bio-Rad Laboratories Pty Ltd, Gladesville, NSW, AUS) and lysates were then diluted with an equal volume in 2x Laemmli buffer containing 10% B-mercaptoethanol.

##### Western blotting

For each protein of interest, a signal linearity test was conducted to determine the ideal loading amount. Muscle lysates were then loaded in equal amounts (10 to 20 μg) and separated by electrophoresis for 1.5 to 2.5 h at 100 V using pre-cast stain-free SDS-PAGE gels (4-20%). Once resolved, the gels were wet transferred onto LF PVDF membranes using a Turbo Transfer system (Bio-rad Laboratories Pty Ltd, Gladesville, NSW, AUS). Membranes were blocked at room temperature for 1 h using 5% skim milk or 5% bovine serum albumin (BSA) in tris buffer saline (TBS) 0.1% tween-20 (TBS-T). After 3 × 5-min washes in TBS-T, membranes were incubated overnight at 4 °C with gentle agitation in primary antibody solutions (1:1000 antibody in 5% BSA, plus 0.02% Na Azide). The antibody for LC3B was purchased from Cell Signalling (#3868S) and the antibody for p62 from Abcam (#ab56416). The following morning, membranes were washed 3 × 5-min in TBS-T and subsequently incubated under gentle agitation at room temperature with the appropriate host species-specific secondary antibody for 60-90 min in 1-5% skim milk in TBS-T. Membranes were washed again for 3 × 5-min in TBS-T before being immersed for 5 min under gentle agitation at room temperature in Clarity ECL detection substrate (Bio-rad Laboratories Pty Ltd, Gladesville, NSW, AUS). Protein bands were visualised using a Bio-Rad ChemiDoc imaging system and band densities were determined using Bio-Rad ImageLab software (Bio-Rad Laboratories Pty Ltd, Gladesville, NSW, AUS). All samples for each participant were loaded on the same gel, along with different concentrations of a mixed-homogenate internal standard (IS), and a calibration curve plotted of density against protein amount. From the subsequent linear regression equation, protein abundance was calculated from the measured band intensity for each lane on the gel. Total protein content of each lane was obtained from the stain-free image of the membrane and was used for normalisation of the results.

##### *Ex vivo* autophagy flux assay

The following protocol was adapted from previous studies performing ex vivo autophagy flux in rodents (19, 25). Upon collection of the skeletal muscle sample, two small pieces (~ 10 mg) were placed in 3 mL of oxygenated DMEM CO_2_ independent media (ThermoFisher #18045088) at 37 °C. The tissues were then incubated with continuous oxygenation for 1 h with (‘treated’ sample, with inhibitors), or without (‘untreated’ sample), 60 μL of NH_4_Cl (20 μL·mL^−1^; 40 mM; Sigma Aldrich #S7653) and 30 μL Leupeptin (10 μL·mL^−1^; 100 uM; Sigma Aldrich #L2884). Upon completion of a 1-h incubation, samples were snap-frozen and stored at −80 °C until further analysis. Autophagy flux (Net LC3B-II flux) was obtained by subtraction of the densitometric value of LC3B-II from treated compared to the untreated sample.

#### Statistical analysis

All values are reported as mean ± standard deviation (SD). All statistical analyses were carried out on the raw values normalised to the total protein loading and calibration curve. For Study 1, one-way repeated-measures of ANOVA with Holm-Sidak post-hoc were used. For Study 2, two-way repeated measures of ANOVA were used, and main effects and interactions were further analysed using Holm-Sidak post-hoc tests. For Study 3, a two-tailed paired student’s t-test was used. For Study 4, a one-way ANOVA with Holm-Sidak post-hoc tests were utilised. Effect sizes (ES) were quantified and defined as: small (0.2), moderate (0.5), large (0.8), and very large (1.3). Statistical significance was set at p < 0.05 for all analyses. GraphPad Prism 8.3 software was used for the statistical analysis.

## Results

### Study 1 - Exercise-induced changes in LC3B and p62 protein changes in soleus muscle of Wistar rat

There was a main effect of time for LC3B-I and LC3B-II protein levels (both p = 0.01), as well as the LC3B-II/I ratio (p = 0.0003). Compared to REST, there was a significant increase in LC3B-I protein level at + 0 h (+ 109 ± 103%; ES = 1.1; p = 0.017; Figure 2.A) and at + 3 h (+ 82 ± 62%; ES = 1.1; p = 0.04; Figure 2.A). Compared to REST, LC3B-II protein level did not significantly change at + 0 h (− 20 ± 46%; ES = − 0.32; p = 0.63; Figure 2.B), but significantly increased at + 3 h (+ 97 ± 102 %; ES = 0.95; p = 0.04; Figure 2.B), and from + 0 h to + 3 h (+ 159 ± 129%; ES = 1.2; p = 0.02; Figure 2.B). There was a significant decrease in the LC3B-II/I ratio immediately (+ 0 h) after exercise (− 65 ± 12 %; ES = − 1.5; p = 0.001; Figure 2.C), followed by a significant increase from + 0 h to + 3 h (+ 164 ± 98%; ES = 1.5; p = 0.001; Figure 2.C). Protein level of p62 did not significantly change at any time point (p > 0.05; Figure 2.D).

**Figure 1.**
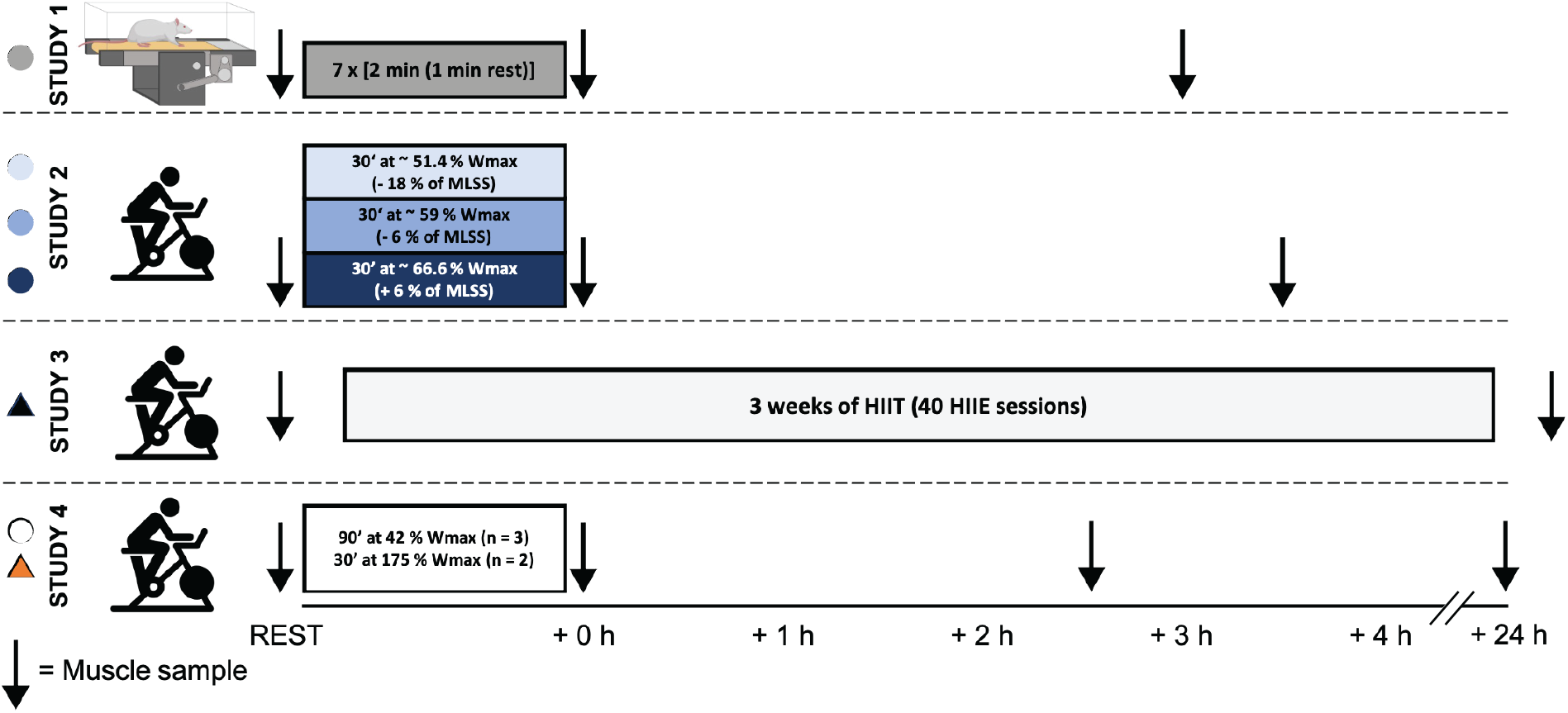
Schematic representation of the four different experimental studies with the time-course of the muscle samples collected. MLSS = Maximal lactate steady state; HIIT = High-intensity interval training; HIIE = High-intensity interval exercise; Ẇ_max_ = maximal aerobic power determined from a graded exercise test. Created with BioRender.com

**Figure 2.**
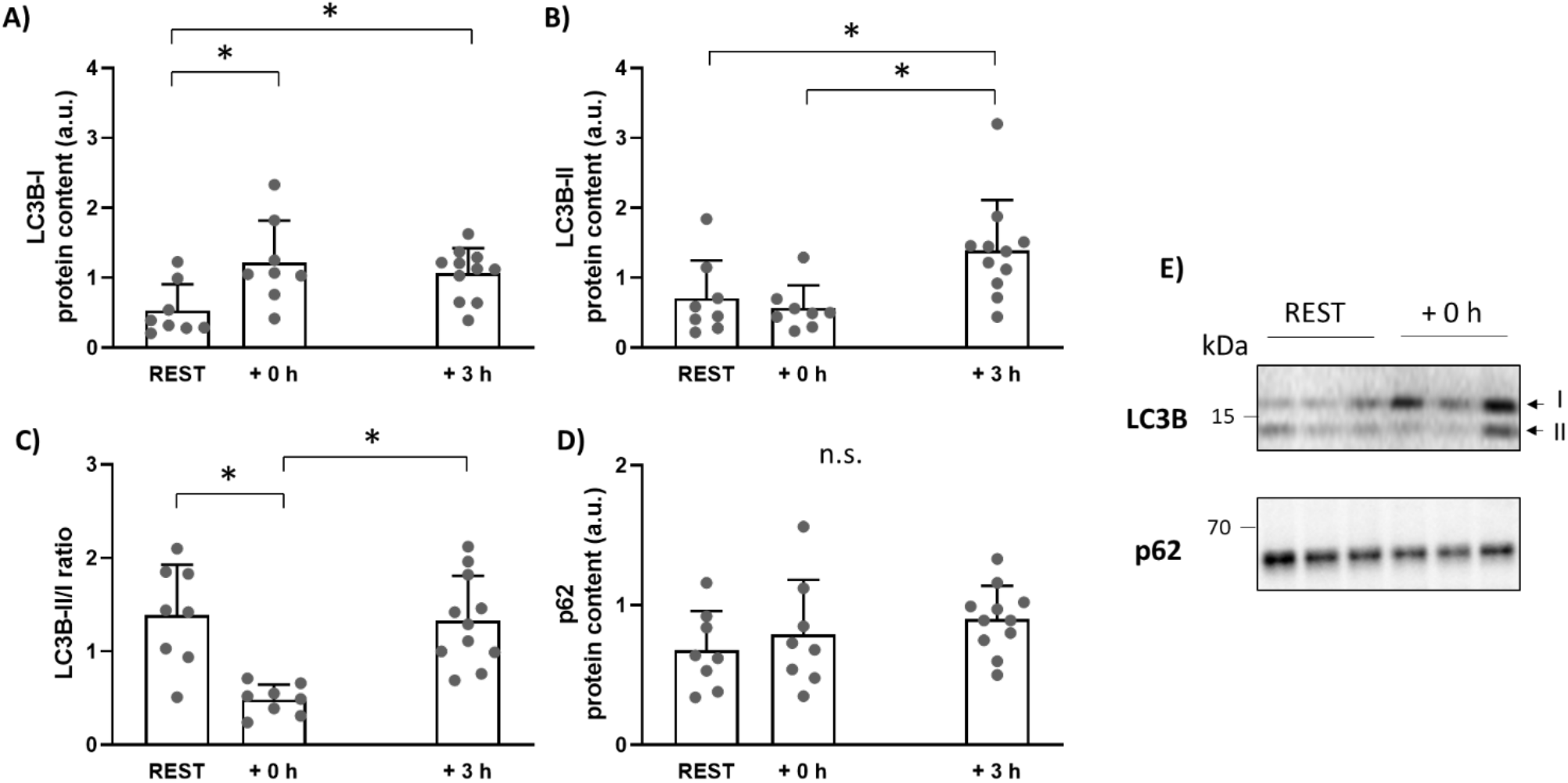
Effects of exercise on A) LC3B-I and; B) LC3B-II protein levels; C) the LC3BII/I ratio and; D) p62 protein levels in the soleus muscle of Wistar rats; E) representative blots of LC3B and p62 protein. Data were analysed using one-way ANOVA; * = p < 0.05. Bars are shown as mean + SD; n.s = not significant.

### Study 2 - Effects of exercise intensity on exercise-induced LC3B and p62 protein changes in human skeletal muscle

There was no main or interaction effect for LC3B-I protein levels (p > 0.05; Figure 3.A). There was no time × intensity effect for LC3B-II protein levels (p = 0.85), but there was a main effect of time (p < 0.0001). Compared to REST, there was a significant decrease at + 0 h (− 24 ± 16%; 90 % CI [− 28, −19%]; ES = − 0.82; p = 0.0001; Figure 3.B), but not at + 3 h (+ 6 ± 31%; 90 % CI [− 4, 15%]; ES = 0.01; p > 0.99; Figure 3.B), and a significant increase between + 0 h and + 3 h (+ 40 ± 38%; 90 % CI [28, 51%]; ES = 0.85; p < 0.0001; Figure 3.B). There was no time × intensity effect for LC3B-II/I ratio (p = 0.85), but there was a main effect of time (p < 0.0001). Compared to REST, there was a significant decrease at + 0 h (− 21 ± 17%; 90 % CI [− 26, − 16%]; ES = − 0.65; p = 0.003; Figure 3.C), but no significant difference at + 3 h (+ 10 ± 45 %; 90 % CI [− 4, 24%]; ES = 0.18; p = 0.27; Figure 3.C), and a significant difference between + 0 h and + 3 h (+ 40 ± 45%; 90 % CI [27, 54%]; ES = 0.67; p = 0.0001; Figure 3.C). There was no main or interaction effect for p62 protein levels (all p > 0.05; Figure 3.D).

**Figure 3.**
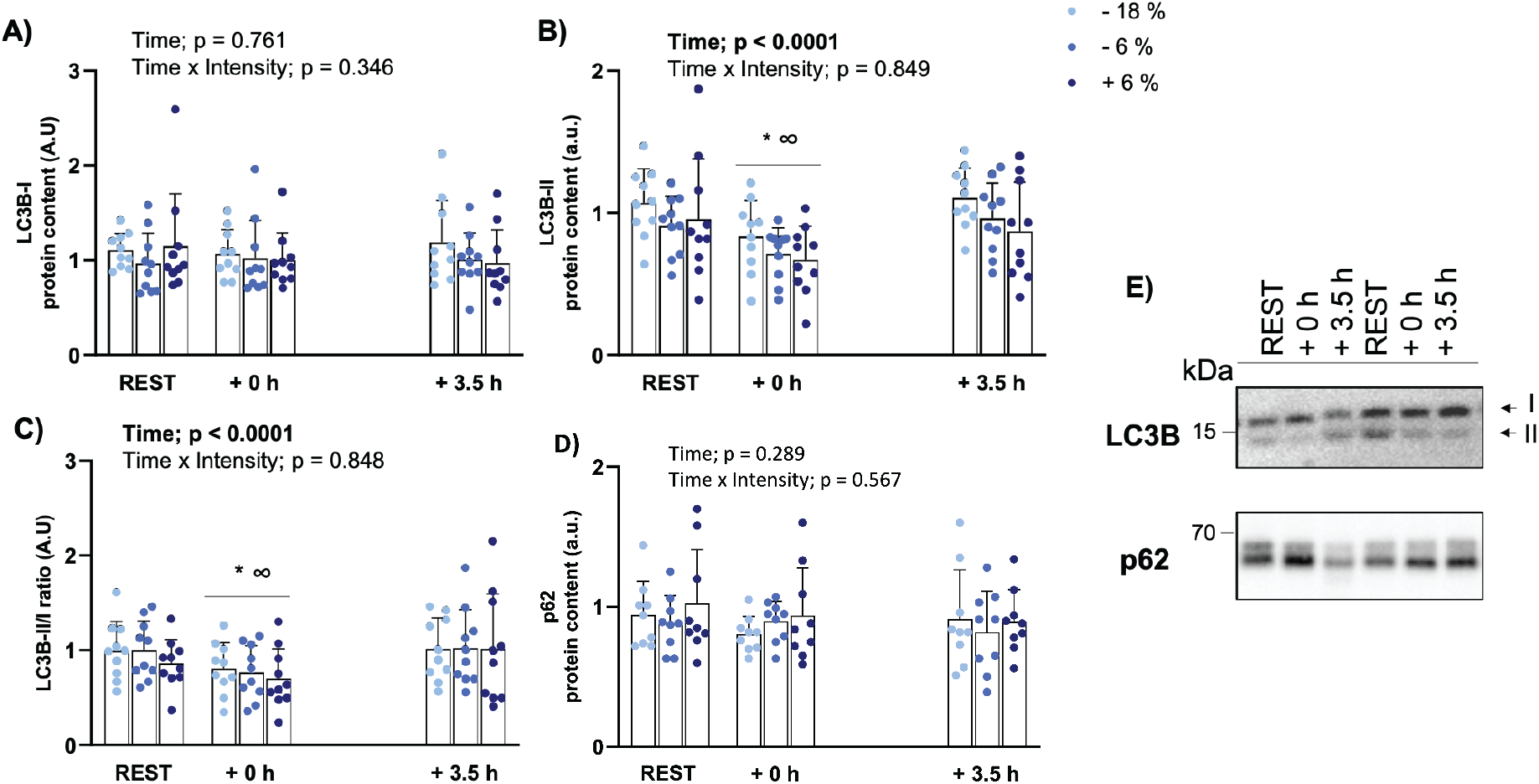
Protein levels of A) LC3B-I and; B) LC3B-II at rest, as well as 0 h and 3.5 h after the end of exercise; C) the LC3BII/I ratio and; D) p62 protein levels at rest, 0 h and 3.5 h after the end of exercise; E) representative blots of LC3B and p62 protein. Participants performed the exercise at three different intensities (−18% = light blue, −6% = normal blue, and +6% = dark blue of the individually determined maximal lactate steady state). n = 9 for p62, n = 10 for LC3B. * = different than REST; ∞ = different from + 3.5 h. Bars shown are mean + SD.

### Study 3 – Effects of high-volume HIIT on resting LC3B and p62 protein levels in human skeletal muscle

Resting levels of LC3B-II protein levels significantly increased from PRE to POST (+ 132 ± 140%; 90 % CI [+ 55, 209%]; ES = 0.85; p = 0.04; Figure 4.A). There was no significant training effect on LC3B-I, LC3B-II/I ratio, and p62 protein levels (all p > 0.05).

**Figure 4.**
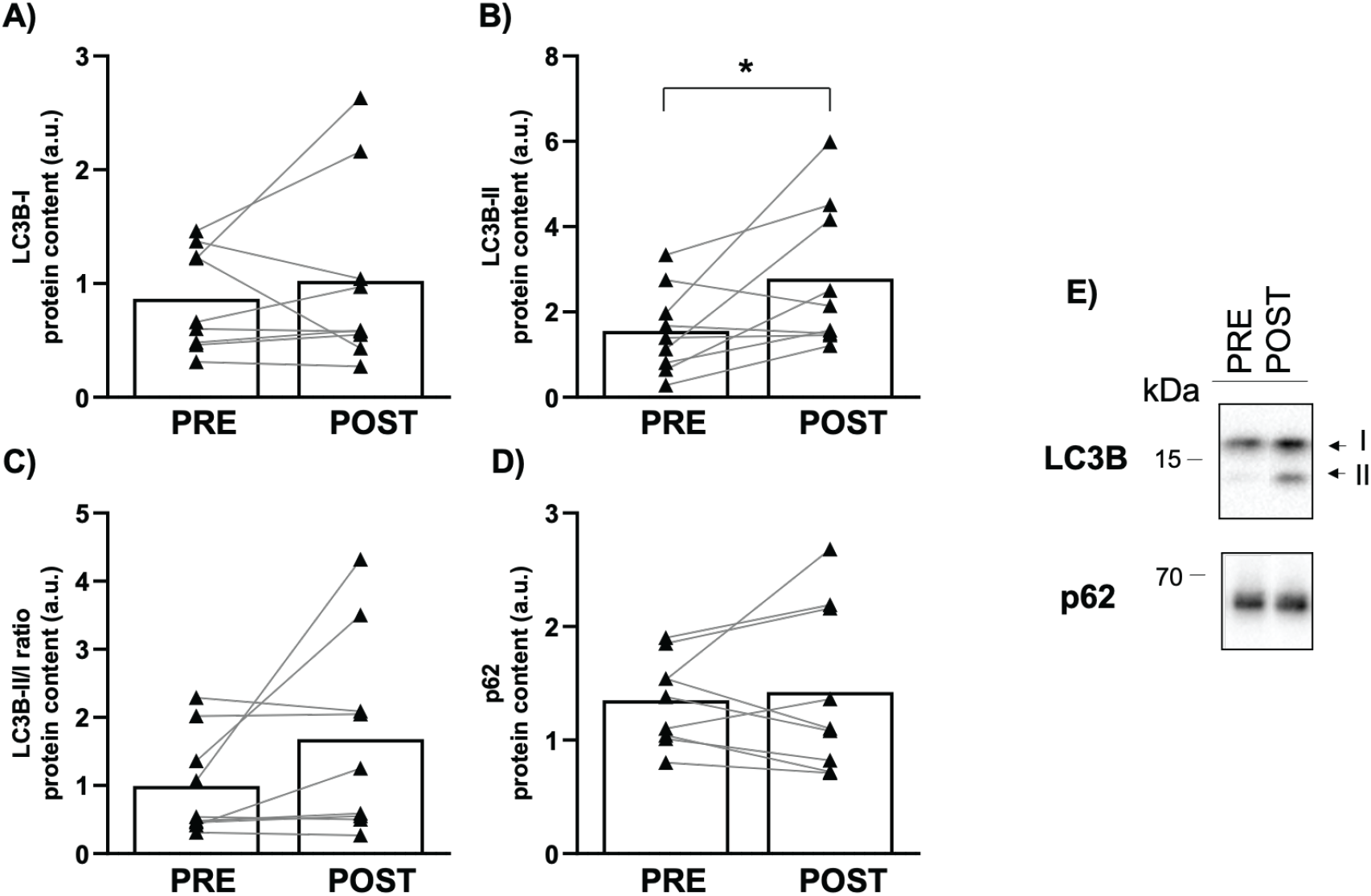
Protein levels of A) LC3B-I and B) LC3B-II in PRE and POST training samples; C) the LC3BII/I ratio and D) p62 protein levels in PRE and POST training samples; E) representative blots of LC3B and p62 proteins. * = significantly different than PRE training sample (p < 0.05). Individual and mean changes are shown.

### Study 4 - Exercise-induced changes in autophagy flux in human skeletal muscle

There was a main effect for LC3B-II protein levels in untreated samples (p = 0.017). Compared to REST there were no significant changes at + 0 h (− 26 ± 23%; 90 % CI [− 43, − 10%]; ES = − 0.77; p = 0.10), at + 2.5 h (+ 23 ± 27%; 90 % CI [3, 43%]; ES = 0.64; p = 0.24), or at + 24 h (+ 8.7 ± 23%; 90% CI [-8, 26%]; ES = 0.26; p = 0.53). However, relative changes from REST to + 0 h between the samples from Study 1 and those in Study 2 were comparable (− 26% vs − 24% respectively; and ES = − 0.77 vs − 0.82 respectively), suggesting a similar exercise-induced response in LC3B-II across experiments in the untreated samples.

For net LC3B-II flux there was no main effect (p = 0.27). Compared to REST, effect size analyses showed a large positive change at + 0 h (+ 117 ± 163%; 90 % CI [− 3, 237%]; ES = 0.82; p = 0.22), at + 2.5 h (+ 113 ± 178%; 90 % CI [− 18, 244%]; ES = 0.88; p = 0.22), and at + 24 h (+ 93 ± 126%; 90% CI [1, 126%]; ES = 0.79; p = 0.22).

## Discussion

Our study shows that: 1) exercise-induced changes in LC3B protein levels differs between rodents and humans; 2) exercise-induced changes in LC3B and p62 protein levels appear to be independent of exercising below or above the MLSS in human skeletal muscle; and 3) the exercise-induced decrease in LC3B-II protein levels observed in humans were not reflective of a decrease in autophagy flux.

The results of the present study showed that the exercise-induced changes in LC3B protein levels differ between rodents and humans. In our rat study, there were increased LC3B-I protein levels 0 to 3 hours following a single endurance exercise session (Figure 2). This was not observed in our human study, in accordance with previous literature (16). An increase of LC3B-I protein levels may stem from increased LC3B mRNA translation. In fact, LC3B mRNA expression has been shown to increase following exercise to exhaustion in mice (10), and an exercise-induced increase in LC3-I protein levels has also been shown in rodents (11). However, it is difficult to compare across studies as not many studies report the changes of LC3-I protein levels.

In the present study, LC3B-II protein levels were unaltered immediately after exercise in rats but were significantly increased 3 hours into the recovery. This is in line with previous research showing that LC3-II is significantly increased in rodents 80 to 180 min from the start of exercise (4, 16, 26). Due to the incomplete information regarding the antibodies used, it was impossible to recapitulate the findings for the LC3 subfamily members used in some of the rodent studies. Future research should address whether the different LC3 subfamily members are similarly modified following exercise in rodents. Our results show that the protein levels of the autophagy receptor p62 remained unchanged at all time points in rats. While this is in contrast to some rodent studies (4, 27), it is in agreement with other findings (10–12, 28). A possible explanation may relate to the duration of the exercise in the different studies, as the only two studies reporting an exercise-induced decrease in p62 protein levels did exercise for at least 110 min (4, 27), and a decrease in p62 was not seen at earlier time points or in the recovery period (27). On the other hand, p62 protein level has been previously shown to be decreased 6 hours into the recovery from both low- and moderate-intensity exercise (12), suggesting a delayed lysosomal degradation of autophagosomes, which may have been missed by most studies including the present study. It is important to mention that other proteins can also act as autophagy receptors (e.g., NBR1, OPTN (29)), and how these are altered by exercise requires further investigation.

In contrast to rodents, the findings from our human study show that, independently of exercising below or above the MLSS, LC3B-II protein levels and the LC3B-II/I ratio were decreased following exercise and returned to baseline 3.5 hours into the recovery (Figure 3). This was in accordance with most human studies (13–16), with one exception (17). A major difference with the study of Brandt, Gunnarsson (17) was the protein analysed. In contrast to the present study and others where an antibody targeting the LC3B subfamily was utilised (13–16), Brandt, Gunnarsson (17) used an antibody targeting a combination of LC3A and LC3B. Whether the protein levels of the different the LC3 subfamily members are differentially regulated following exercise remains to be elucidated. Interestingly, a proteomic analysis of human skeletal muscle studies only detected LC3A (30), which may indicate a greater protein abundance of LC3A when compared to the other subfamily members in skeletal muscle. The present findings also show that LC3B-I protein levels were not altered following exercise, consistent with previous studies in humans (13, 15). The unchanged LC3B-I protein levels could be due to unchanged mRNA expression of LC3B or rapid conjugation of LC3-I into LC3-II and increased autophagosome degradation. The finding of unchanged p62 protein levels following exercise, independent of exercise intensity, were in accordance with most studies (13, 15–17). Furthermore, the previously reported role of exercise intensity on p62 protein changes (14) may not be due to exercise intensity differences between protocols, but possibly related to other factors such as total work performed.

Our data demonstrated that following three weeks of high-volume HIIT there was an increase in basal LC3B-II protein level, suggesting an increase in autophagosome content (Figure 4). To the best of our knowledge, this is the first study suggesting that HIIT can lead to increased autophagosome content but our findings are in line with the results of a previous study where three weeks of one-legged knee extensor training led to an increase in LC3B-II protein levels (16). However, others have not shown any effect of endurance training on LC3A/B-II protein levels, despite an increase in LC3A/B-I protein levels (17). Whether training volume or intensity are more important for the training-induced changes in LC3B-II requires further research.

A limitation of human studies to date is the use of LC3B-II and p62 protein levels to infer changes in autophagy flux. This has led to the idea that a decrease in LC3B-II protein levels following exercise could be reflective of a temporary decrease in autophagy flux (13). In the present study, a protocol adapted from a rodent study (19) was used to examine, for the first time, the effects of exercise on *ex vivo* autophagy flux in human skeletal muscle. The results showed that autophagy flux (measured as net LC3B-II flux) did not decrease immediately after 0, 2.5, or even 24 hours after exercise (Figure 5.A). Although limited by the low number of participants, the effect size analyses suggested a moderate-to-large increase in *ex vivo* autophagy flux following exercise (+ 93-117%; ES = 0.79-0.88). Our findings in humans were in agreement with those from a rodent study showing a similar exercise-induced fold-change in autophagy flux (10). These findings would suggest that exercise-induced autophagy flux is similar between rodents and humans (Figure 5.B), despite a different exercise-induced LC3B protein regulation. The use of an *ex vivo* autophagy flux assay in future human studies will allow researchers to overcome the limitation of solely relying on static protein markers. Future research should interrogate the autophagy flux response to different stimuli (e.g., inactivity) in skeletal muscle and with larger sample sizes.

**Figure 5.**
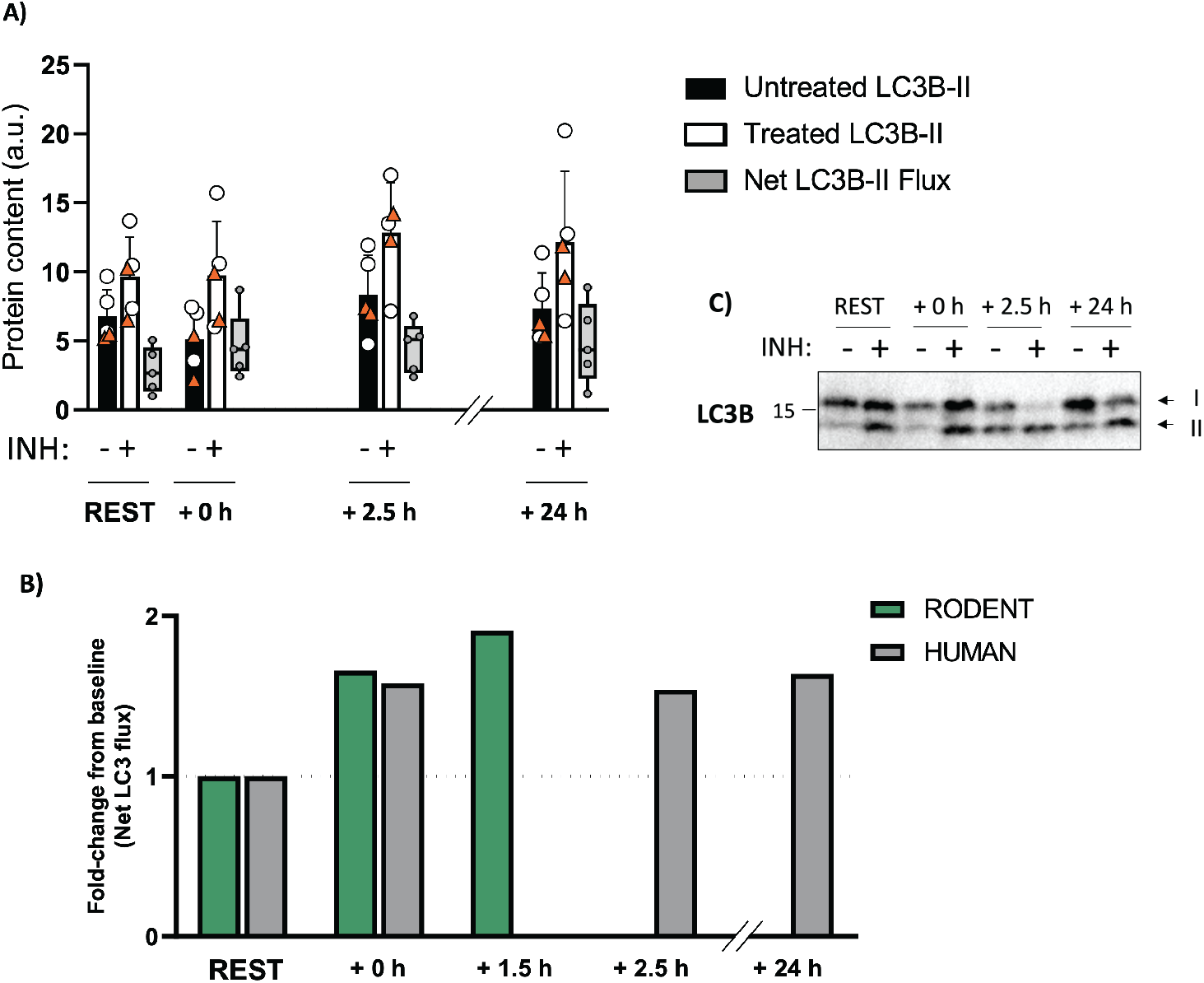
A) LC3B-II protein levels from untreated (black bars) and treated samples (white bars), and the net LC3B-II flux (in grey; calculated by subtracting untreated LC3B-II protein levels from treated sample). C) Representative blot. Orange triangles represent participants performing SIE (n = 2); white circles represent participants performing MICE (n = 3). B) Fold-change following exercise in the net LC3-II flux from the present study (LC3B-II, human) and a published rodent study (LC3-II, adapted from (10)). Bars for the treated and untreated samples display the mean + SD. Individual data points along with box and whisker plots are shown for net LC3B-II flux. INH = inhibitors NH_4_Cl (40 mM) and Leupeptin (100 μM) added to the treated sample.

The main limitation of the present study is the low sample size and the two different exercises utilised in our *ex vivo* autophagy flux experiments. Nonetheless, our findings highlight the value of using this assay in human skeletal muscle studies. Furthermore, our study was limited to LC3B, whereas the role of other ATG8 family members in exercise and skeletal muscle autophagy remains unexplored.

In conclusion, the results of the current study showed that exercise-induced LC3B protein changes differ between rodents and humans. This indicates caution must be taken when extrapolating autophagy protein results from rodents to humans. Furthermore, findings from the current study show that a reduction in LC3B-II protein levels following exercise in humans was consistent across exercise intensities but was not indicative of a decrease in autophagy flux. This suggests that studies should avoid looking at ‘static’ levels of LC3 protein levels and include autophagy flux assays to provide a more valid assessment of dynamic changes in autophagy with exercise.

## Abbreviations

LC3: Microtubule-associated proteins 1A/1B light chain 3
p62: ubiquitin-binding protein p62
INH: inhibitors
MLSS: maximal lactate steady state
HIIE: high-intensity interval exercise
HIIT: high-intensity interval training
MICE: moderate-intensity continuous exercise
SIE: sprint-interval exercise
GXT: graded exercise test
V̇O_2peak_: peak oxygen uptake
V̇O_2max_: maximal oxygen uptake W_LT_, power at the lactate threshold
Ẇ_max_: maximal aerobic power

## Acknowledgements

The authors would like to thank all the participants for volunteering to take part in these studies.

## Conflict of Interest statement

All authors involved in this research declare no conflict of interest.

## Funding

This work was supported by an ARC grant to D.J.B (DP140104165).

## Author contributions

J.B, C.G, N.A.J, A.J.G, M.L, and D.J.B designed the research. J.B, C.G, N.A.J, A.J.G, E.P, T.J, and A.G performed the research. J.B, M.L, and D.J.B wrote the manuscript. J.B, C.G, N.A.J, A.J.G, E.P, T.J, A.G, M.L, and D.J.B revised and approved the manuscript.

## Notes

### Competing Interest Statement

The authors have declared no competing interest.

## References

1. Choi, A. M., Ryter, S. W., and Levine, B. (2013) Autophagy in human health and disease. The New England journal of medicine 368, 651–662

2. Lo Verso, F., Carnio, S., Vainshtein, A., and Sandri, M. (2014) Autophagy is not required to sustain exercise and PRKAA1/AMPK activity but is important to prevent mitochondrial damage during physical activity. Autophagy 10, 1883–1894

3. Call, J. A., Wilson, R. J., Laker, R. C., Zhang, M., Kundu, M., and Yan, Z. (2017) Ulk1-mediated autophagy plays an essential role in mitochondrial remodeling and functional regeneration of skeletal muscle. American journal of physiology. Cell physiology 312, C724–c732

4. He, C., Bassik, M. C., Moresi, V., Sun, K., Wei, Y., Zou, Z., An, Z., Loh, J., Fisher, J., Sun, Q., Korsmeyer, S., Packer, M., May, H. I., Hill, J. A., Virgin, H. W., Gilpin, C., Xiao, G., Bassel-Duby, R., Scherer, P. E., and Levine, B. (2012) Exercise-induced BCL2-regulated autophagy is required for muscle glucose homeostasis. Nature 481, 511–515

5. Lira, V. A., Okutsu, M., Zhang, M., Greene, N. P., Laker, R. C., Breen, D. S., Hoehn, K. L., and Yan, Z. (2013) Autophagy is required for exercise training-induced skeletal muscle adaptation and improvement of physical performance. FASEB journal: official publication of the Federation of American Societies for Experimental Biology 27, 4184–4193

6. Melia, T. J., Lystad, A. H., and Simonsen, A. (2020) Autophagosome biogenesis: From membrane growth to closure. The Journal of cell biology 219

7. Slobodkin, M. R., and Elazar, Z. (2013) The Atg8 family: multifunctional ubiquitin-like key regulators of autophagy. Essays in biochemistry 55, 51–64

8. Nguyen, T. N., Padman, B. S., Usher, J., Oorschot, V., Ramm, G., and Lazarou, M. (2016) Atg8 family LC3/GABARAP proteins are crucial for autophagosome–lysosome fusion but not autophagosome formation during PINK1/Parkin mitophagy and starvation. Journal of Cell Biology 215, 857–874

9. Tanida, I., Ueno, T., and Kominami, E. (2004) LC3 conjugation system in mammalian autophagy. The international journal of biochemistry & cell biology 36, 2503–2518

10. Vainshtein, A., Tryon, L. D., Pauly, M., and Hood, D. A. (2015) Role of PGC-1α during acute exercise-induced autophagy and mitophagy in skeletal muscle. American journal of physiology. Cell physiology 308, C710–719

11. Zhang, D., Lee, J. H., Kwak, S. E., Shin, H. E., Zhang, Y., Moon, H. Y., Shin, D. M., Seong, J. K., Tang, L., and Song, W. (2019) Effect of a Single Bout of Exercise on Autophagy Regulation in Skeletal Muscle of High-Fat High-Sucrose Diet-Fed Mice. Journal of Obesity & Metabolic Syndrome 28, 175–185

12. Brandt, N., Dethlefsen, M. M., Bangsbo, J., and Pilegaard, H. (2017) PGC-1α and exercise intensity dependent adaptations in mouse skeletal muscle. PloS one 12, e0185993

13. Kruse, R., Pedersen, A. J., Kristensen, J. M., Petersson, S. J., Wojtaszewski, J. F., and Hojlund, K. (2017) Intact initiation of autophagy and mitochondrial fission by acute exercise in skeletal muscle of patients with Type 2 diabetes. Clinical science (London, England: 1979) 131, 37–47

14. Schwalm, C., Jamart, C., Benoit, N., Naslain, D., Premont, C., Prevet, J., Van Thienen, R., Deldicque, L., and Francaux, M. (2015) Activation of autophagy in human skeletal muscle is dependent on exercise intensity and AMPK activation. FASEB journal: official publication of the Federation of American Societies for Experimental Biology 29, 3515–3526

15. Moller, A. B., Vendelbo, M. H., Christensen, B., Clasen, B. F., Bak, A. M., Jorgensen, J. O., Moller, N., and Jessen, N. (2015) Physical exercise increases autophagic signaling through ULK1 in human skeletal muscle. Journal of applied physiology (Bethesda, Md.: 1985) 118, 971–979

16. Fritzen, A. M., Madsen, A. B., Kleinert, M., Treebak, J. T., Lundsgaard, A. M., Jensen, T. E., Richter, E. A., Wojtaszewski, J., Kiens, B., and Frosig, C. (2016) Regulation of autophagy in human skeletal muscle: effects of exercise, exercise training and insulin stimulation. The Journal of physiology 594, 745–761

17. Brandt, N., Gunnarsson, T. P., Bangsbo, J., and Pilegaard, H. (2018) Exercise and exercise training-induced increase in autophagy markers in human skeletal muscle. Physiological reports 6, e13651–e13651

18. Yoshii, S. R., and Mizushima, N. (2017) Monitoring and Measuring Autophagy. Int J Mol Sci 18, 1865

19. Martinez-Lopez, N., Tarabra, E., Toledo, M., Garcia-Macia, M., Sahu, S., Coletto, L., Batista-Gonzalez, A., Barzilai, N., Pessin, J. E., Schwartz, G. J., Kersten, S., and Singh, R. (2017) System-wide Benefits of Intermeal Fasting by Autophagy. Cell metabolism 26, 856–871.e855

20. Jamnick, N. A., Botella, J., Pyne, D. B., and Bishop, D. J. (2018) Manipulating graded exercise test variables affects the validity of the lactate threshold and [Formula: see text]. PloS one 13, e0199794

21. Billat, V. L., Sirvent, P., Py, G., Koralsztein, J. P., and Mercier, J. (2003) The concept of maximal lactate steady state: a bridge between biochemistry, physiology and sport science. Sports medicine (Auckland, N.Z.) 33, 407–426

22. Granata, C., Oliveira, R. S. F., Little, J. P., and Bishop, D. J. (2020) Forty high-intensity interval training sessions blunt exercise-induced changes in the nuclear protein content of PGC-1α and p53 in human skeletal muscle. American journal of physiology. Endocrinology and metabolism 318, E224–e236

23. Granata, C., Oliveira, R. S. F., Little, J. P., Renner, K., and Bishop, D. J. (2016) Mitochondrial adaptations to high-volume exercise training are rapidly reversed after a reduction in training volume in human skeletal muscle. FASEB journal: official publication of the Federation of American Societies for Experimental Biology 30, 3413–3423

24. Granata, C., Oliveira, R. S., Little, J. P., Renner, K., and Bishop, D. J. (2016) Training intensity modulates changes in PGC-1alpha and p53 protein content and mitochondrial respiration, but not markers of mitochondrial content in human skeletal muscle. FASEB journal: official publication of the Federation of American Societies for Experimental Biology 30, 959–970

25. Yamada, E., and Singh, R. (2012) Mapping autophagy on to your metabolic radar. Diabetes 61, 272–280

26. Jamart, C., Naslain, D., Gilson, H., and Francaux, M. (2013) Higher activation of autophagy in skeletal muscle of mice during endurance exercise in the fasted state. American journal of physiology. Endocrinology and metabolism 305, E964–974

27. Pagano, A. F., Py, G., Bernardi, H., Candau, R. B., and Sanchez, A. M. (2014) Autophagy and protein turnover signaling in slow-twitch muscle during exercise. Medicine and science in sports and exercise 46, 1314–1325

28. Fritzen, A. M., Frøsig, C., Jeppesen, J., Jensen, T. E., Lundsgaard, A. M., Serup, A. K., Schjerling, P., Proud, C. G., Richter, E. A., and Kiens, B. (2016) Role of AMPK in regulation of LC3 lipidation as a marker of autophagy in skeletal muscle. Cellular signalling 28, 663–674

29. Behrends, C., and Fulda, S. (2012) Receptor proteins in selective autophagy. Int J Cell Biol 2012, 673290–673290

30. Gonzalez-Freire, M., Semba, R. D., Ubaida-Mohien, C., Fabbri, E., Scalzo, P., Højlund, K., Dufresne, C., Lyashkov, A., and Ferrucci, L. (2017) The Human Skeletal Muscle Proteome Project: a reappraisal of the current literature. Journal of Cachexia, Sarcopenia and Muscle 8, 5–18

